# Axon demyelination and degeneration in a zebrafish *spastizin* model of hereditary spastic paraplegia

**DOI:** 10.1101/2024.04.15.589631

**Authors:** Vranda Garg, Luisa Heyer, Torben Ruhwedel, Selina André, Gudrun Kracht, Patricia Scholz, Till Ischebeck, Hauke B. Werner, Christian Dullin, Jacob Engelmann, Wiebke Möbius, Roland Dosch, Martin C. Göpfert, Bart R.H. Geurten

## Abstract

Hereditary spastic paraplegias (HSPs) are a diverse set of neurological disorders characterized by progressive spasticity and weakness in the lower limbs caused by damage to the axons of the corticospinal tract. More than 88 genetic mutations have been associated with HSP, yet the mechanisms underlying these disorders are little understood. We studied the pathogenesis of one form of HSP known as spastic paraplegia 15 (SPG15). This disorder is caused by mutations in the *ZFYVE26* gene, which codes for a protein called SPASTIZIN. We show that, in zebrafish, the significant reduction of Spastizin caused degeneration of Mauthner (M)-cells. M-cell degeneration is associated with axon demyelination in the spinal cord and impaired locomotion in the *spastizin* mutants. Our findings reveal that the mutation not only compromises axonal integrity but also affects the structural molecules of the myelin sheath, laying the foundation for degeneration and advancing our understanding of the intricate mechanisms underlying HSPs.

## 3 Introduction

Hereditary spastic paraplegias (HSPs) are a group of debilitating neurological disorders passed down through families. These conditions are characterised by progressive weakness and spasticity in the lower limbs, making it difficult for affected individuals to move freely and to perform daily tasks. The cause of these symptoms is the distal neuropathy of the longest corticospinal tract axons, which control lower limb motility (Blackstone, 2012, 2018, Fink, 2006, 2014, Finsterer et al., 2012, Harding, 1993, 1984, Klebe et al., 2015, Tesson et al., 2015). To date, more than 88 different genes and 100 distinct spastic gait disease loci have been identifed to cause HSP (D’Amore et al., 2018, Elsayed et al., 2021, Galatolo et al., 2018, Giudice et al., 2014, Hensiek et al., 2015, Parodi et al., 2018). The proteins encoded by these genes are involved in a variety of cellular processes (Blackstone, 2012, 2018, Fink, 2014, Giudice et al., 2014). In the case of SPG15, the affected protein Spastizin is mostly involved in endosomal trafficking (Khundadze et al., 2013), autophagy (Vantaggiato et al., 2013, 2019), lysosomal biogenesis (Chang et al., 2014) and cytokinesis (Sagona et al., 2010). The 285 kDa protein is widely distributed inside cells and co-localizes with a variety of organelles including endosomes, microtubules, endoplasmic reticulum (ER), mid-body, and vesicles involved in protein trafficking (Hanein et al., 2008, Murmu et al., 2011). Spastizin interacts with Spatacsin and Beclin-1 (Sagona et al., 2011, Słabicki et al., 2010) and is essential for the proper development and outgrowth of motor neuron axons (Martin et al., 2012).

To date, relatively few studies have addressed the pathophysiology of SPG15, with most of them using mouse models. For example, Khundadze et al. (2013) created *zfyve26* knockout mice, that developed late-onset spastic paraplegia and cerebellar ataxia. Judging from this mutant, endolysosomal dysfunction might cause the degeneration of cortical motor neurons and Purkinje cells in the cerebellum. Despite considerable knowledge about the functions of Spastizin in different cellular processes, its role in the pathogenesis of HSP remains poorly understood.

In this study, we generated a model for the SPG15 in zebrafish and found that the significant reduction of Spastizin caused a severe demyelination and degeneration of the spinal cord motor axons that leads to locomotion defects in *spastizin* mutants. Moreover, this truncating mutation affects the activity of the large motor neurons including the M-cells which are believed to be the functional reminiscent of lower limb motor neurons (Eaton et al., 1977, Faber and Korn, 1978, Korn and Faber, 2005). Our work characterises the crucial role of Spastizin in maintaining neuronal integrity, emphasising its profound impact on motor neuron function and underscoring its significance in the complex pathogenesis of HSP.

## 4 Materials and methods

### 4.1 Zebrafish husbandry

The zebrafish were maintained according to the guidelines provided by the Westerfeld zebrafish book (Westerfield, 2000) and EuFishBioMed/Felasa (Aleström et al., 2020), in compliance with the regulations of the Georg-August University, Goettingen, and Bielefeld University, Bielefeld, Germany. Preliminary studies in our lab indicated a late onset of HSP symptoms in *spastizin* mutants, therefore all the experiments were conducted on adult fish between 10-15 months of age.

### 4.2 Ethical approval declarations

The zebrafish experiments were approved by the Lower Saxony State Office for Consumer Protection and Food Safety (AZ14/1681/Dosch) and were performed according to EU directive 2010/63/EU.

### 4.3 *spastizin* mutant generation

The target for *zfyve26* was: 5’CTCACCGCGCTGCAGACA3’. In vitro transcribed guide RNA and Cas9 RNA were co-injected in +/- embryos produced by crossing homozygous males from another *spastizin* mutant line, known as *souffle^p96re^*, with wild-type AB*TLF females. The *souffle^p96re^* mutants carry a point mutation in the splice donor site, producing a shortened 2270 amino acid protein (Kanagaraj et al., 2014). The injected embryos were developed to adulthood, and the success of the gene editing method was assessed using the T7 Endonuclease 1 (T7E1) assay kit (Invitrogen, Germany). Positive individuals from the resulting F1 generation were then crossed with wild-type fish to produce the F2 generation which were screened again with T7E1 assay. The positive individuals in the F2 generation were crossed with each other to produce the F3 generation. In this generation, two males and four females were found to have a 4 base pair deletion in one of the alleles, resulting in a pre-mature stop codon. These heterozygous individuals were used for breeding and creating a new *spastizin* mutant line which was named as *zfyve26^ge1^*.

### 4.4 Genotyping

The melting curve analysis method was used to identify heterozygous and homozygous *spastizin* mutants, using wild-type fish as a positive control [Supplementary Figure 2]. First, tail fin clips were cut and frozen at -80*^◦^*C for 30 minutes. For extracting DNA, 100 µl lysis buffer containing 10 µl 0.1M Tris pH 8.5 (Carl Roth, Germany), 5 µl proteinase K (20mg/ml) (Merck (Sigma-Aldrich), Germany), and 85 µl distilled water was added to them and the mixture was incubated at 55*^◦^*C for 4 hours. The proteinase K was inactivated by heating the mixture at 95*^◦^*C for 10 minutes. A master mix containing 4.2 µl distilled water, 0.4 µM of each primer (Merck (Sigma-Aldrich), Germany) from a working concentration of 10 µM, 5 µl precision melt supermix (Bio-Rad, Germany) and 1 µl DNA was used for real-time PCR. The mixture was subjected to the following PCR conditions: 95*^◦^*C for 3 min; 95*^◦^*C for 5 sec; 65*^◦^*C for 10 sec; repeat step 2-3 39 more times; 95*^◦^*C for 60sec; 65*^◦^*C for 60 sec in the presence of dsDNA intercalating dye SYBR green. Then a melt curve reaction was performed on the rehybridized DNA, by heating it from 65-95*^◦^*C with 0.1*^◦^*C increase every 5 seconds with plate read, and later analysed using Bio-rad CFX Maestro software. The following primer pair was used for the genotyping: Forward 5’-TCCAGGCGGGAGCTCTTC-3’ and Reverse 5’-ATCTTCTGCACGCGCCTC-3’.

### 4.5 Western blot

Brain samples from all the three genotypes were homogenized in radio-immunoprecipitation assay (RIPA) buffer (0.1% sodium dodecyl sulfate (SDS) (Serva GmbH, Germany), 1% Nonidet P-40 (NP40) (Merck (Sigma-Aldrich), Germany), 1% sodium deoxycholate (Merck (Sigma-Aldrich), Germany), 150 mM sodium chloride (NaCl) (Carl Roth, Germany), 50 mM Tris (hydroxymethyl) aminomethane (THAM) hydrochloride (Tris-HCl) (pH = 7.2-7.5) (Carl Roth, Germany), Halt Protease Inhibitor Cocktail (100X) (Thermo Fisher Scientific, Germany)) and centrifuged at 10,000xg for 10 min at 4*^◦^*C. The supernatant was collected and Bradford assay (Bradford, 1976) was conducted to determine the protein concentration. 50 µg lysate from each sample type was separated on a 7% SDS-polyacrylamide gel initially at 30mA until the samples reach the separating gel and then at 40mA, total approximately for 2 hours. The samples on the gel were then electrotransferred to the polyvinylidene difluoride (PVDF) membrane (Bio-Rad, Germany) at 40V for 24 hours in blotting buffer (5% MeOH, 0.05% SDS). For the overnight electrotransfer, the whole electrophoresis set up was placed in a cold room at 4*^◦^*C and the electrophoresis apparatus itself was placed an ice box. The membrane was blocked with 4% powdered milk (Carl Roth, Germany) in phosphate buffered saline (PBS) (ChemSolute, Germany) + 0.1% Tween (AppliChem, Germany) (PBST) for 30 min and incubated with anti-Spastizin (1:300, Biogenes, Germany) (Kanagaraj et al., 2014) or anti-Spastizin (1:200, Invitrogen, Germany) antibody in 3% Bovine serum albumin (BSA) (Carl Roth, Germany) in PBST + 0.02% sodium azide (Carl Roth, Germany), overnight at 4*^◦^*C. Secondary antibody, anti-rabbit horseradish peroxidase (HRP) (1:5000, Merck (Sigma-Aldrich), Germany) was used in blocking solution for 30 min at room temperature. iBright CL1000 (Invitrogen, Germany) was used for capturing and NIH ImageJ/FIJI (Schindelin et al., 2012) was used for analysing images.

A separate gel was used for immunoblotting with anti-*β*-Tubulin (1:1000, DSHB, USA) antibody, because while separating the bands with high molecular weight more clearly, the bands with low molecular weight were already leaked out. All the conditions and procedure was kept same as for Spastizin except the electrotransfer. Samples on the gel were electrotransferred to a nitrocellulose membrane for 1.5 h at 180mA in blotting buffer without additional methanol and SDS. Secondary antibody, anti-mouse horseradish peroxidase (HRP) (1:5000, Merck Sigma-Aldrich), Germany) was used.

### 4.6 Immunohistochemistry

The spinal cord was dissected and divided into four equal parts. The parts were fixed in 4% paraformaldehyde (PFA) (Carl Roth, Germany) in 0.1 M phosphate (PO_4_) buffer (pH = 7.4) and embedded in Albumin-Gelatin. 50 µm thick cross-sections of spinal cord were cut using a Leica VT1000P vibrating blade microtome (Leica, Germany). Sections were blocked with 5% normal goat serum (NGS) (Jackson ImmunoResearch, UK) + 0.25% BSA in PBS/1% Triton X-100 (Merck (Sigma-Aldrich), Germany) for 2h at room temperature. After blocking, the sections were incubated with anti-Spastizin primary antibody (1:300, Biogenes, Berlin, Germany) in blocking solution and incubated at 4*^◦^*C overnight on a stirrer. The following day, the sections were washed six times with PBS-1% Triton X-100 for 10 minutes each. The sections were then incubated with the secondary antibody donkey anti-rabbit AF 488 (1:300, Invitrogen, Germany) or donkey anti-rabbit Cyanine Dye 3 (Cy3) (1:300, Jackson ImmunoResearch, UK) in PBS/1% Triton X-100 in darkness for 3 h at room temperature. After incubation, the sections were washed six times with PBS for 10 minutes each and once with PBS/1,4-diazabicyclo[2.2.2]octane (DABCO) (Carl Roth, Germany) (1:1). Finally, the sections were mounted on slides with PBS/DABCO and stored in slide boxes at 4*^◦^*C until the confocal imaging was performed. To study axonal projections in the spinal cord, the same procedure was followed but using anti-*β*-Tubulin (1:1000, DSHB, USA) as the primary antibody and Goat anti-Mouse AF 546 (1:300, Invitrogen, Germany) as secondary antibody.

### 4.7 Confocal microscopy

Images were captured using a Leica TCS SP8 confocal microscope (Leica, Germany) at a resolution of 2048×2048 pixels with the 20X objective for Spastizin protein confirmation in the Mauthner axons. For axonal diameter measurement, images were acquired at a resolution of 1024×1024 pixels with the 63X glycerol immersion objective. NIH ImageJ/FIJI (Schindelin et al., 2012) was used for image processing and signal intensity measurement in the Mauthner axons. A Matlab R2012b script (The MathWorks Inc., Natick, MA, USA) was used for measuring the neuronal diameter and numbers.

### 4.8 Locomotion recordings

Zebrafish were filmed from above using a Genie HM1024 high-speed camera (Dalsa Imaging Solutions GmbH, Germany) linked with an Optem Zoom 125C 12.5:1 Micro-Inspection Lens System. The setups were illuminated from below with a light emitting diode (LED) light plate (Lumitronix, Germany) and an aquarium light control (Elektronik-Werkstatt SSF, University of Göttingen, Germany). Videos were recorded for 30 seconds at 200 frames per second (fps). The experiments were conducted in the diurnal rhythm between 10 am and 8 pm.

#### 4.8.1 Unmotivated and motivated trials

To investigate the swimming behaviour, zebrafish were filmed for both cruising and motivated trials using the high-speed camera. The experiments were conducted in a 24.9 x 11.4 cm acrylic glass aquarium with a shallow water depth of 1.6 cm. The recording for cruising began 30 seconds after transferring a fish to the setup tank. The motivated trial started immediately after the cruising recording ended. For the motivated trials, a 474 g metal weight was used, that fell through a plastic tunnel and struck the setup table, creating a mechanical stimulus of 18.7 Newton (N) on the surface. By analysing the footage, the fish’s response to the mechanical stimulus was observed.

#### 4.8.2 Counter current trials

A custom-built 17.2 x 4.4 cm Plexiglas aquarium complete with an installed water pump was constructed to study the impact of the water flow on fish movement. This setup allowed to simulate three distinct current velocities of 172, 240, and 277 ml/s, respectively, which were referred as slow, medium, and fast streams. The fish movement was recorded and analysed in these streams using a high-speed camera to understand how they adapt to different water flow conditions.

#### 4.8.3 Electrophysiology

The experimental set up for measuring the electric field potential from Mauthner neurons was adapted (Issa et al., 2011). A set up tank of 8 X 4 cm was used for the present study. The water jet from the picospritzer pressure pump (Parker Hannifin, USA) was used to evoke the escape response of the observed zebrafish. The signal was amplified 2000 times and band-pass filtered with a pass window of 300 to 500 Hz. Additionally, a Hum Bug (Quest Scientific, USA) was used to filter out the electronic noise of 50 Hz. The recording tank was filled with Milli-Q water to achieve a resistance of 18.2 *<u>^M^</u>*<u>^Ω^</u>. The animal movement was recorded at 936 fps for 5 seconds via the video camera that was triggered by the picospritzer. Electrophysiological signals were recorded with the micro2 1401 (Cambridge Electronic Design, UK) data acquisition (DAQ) system and analysed with Spike2 software (Cambridge Electronic Design, UK).

#### 4.8.4 Behavioural data analysis

Zebrafish locomotion was tracked with Limbless Animal traCkEr (LACE), a MATLAB R2012b script (The MathWorks Inc., Natick, Massachusetts, USA) (Garg et al., 2022).

### 4.9 Extraction and analysis of cholesterol

Whole brain samples dissected from wild-type, heterozygous and homozygous *spastizin* mutant fish were kept in pre-weighed reaction cups and immediately flash frozen in liquid nitrogen and stored at -80*^◦^*C until further processing. The samples were lyophilized, weighed and then ground in a ball mill to make a fine powder. Two-phase extraction was performed by adding 400 µl of methyl-tert-butyl ether (MTBE): methanol (3:1, v/v; all solvents of High-performance liquid chromatography (HPLC)-grade (Thermo Fisher Scientific, Germany)), followed by vortexing and the addition of 200 µl water. 5 µg 17:0 free fatty acid (Merck, Germany) was used as the internal standard. After adding the standard, samples were mixed for 30 min on a rotator. The samples were then centrifuged at maximum speed for 1 min and the upper phase was transferred to a fresh tube and stored at -20*^◦^*C until further processing. For gas chromatography-mass spectrometry (GC-MS) measurements, samples were prepared by evaporating 20 *−* 50 µl of the upper phase under a stream of nitrogen, redissolving in 10 *−* 15 µl anhydrous pyridine (Merck, Germany) and derivatizing with twice the volume of N-methyl-N-trimethylsilyltrifluoroacetamid (MSTFA) (Merck, Germany) to yield trimethylsilylated analytes. An Agilent 7890B gas chromatograph connected to an Agilent 5977N mass-selective detector was used for sample analysis, as described in (Berghoff et al., 2021). Cholesterol was identified in comparison to an external standard. For quantification, the peak area of the mass-to-charge ratios 327 Da/e and 458 Da/e were used for the internal standard and cholesterol, respectively. Quantification was done using the Agilent software GC/MSD MassHunter with MSD ChemStation Data Analysis. Cholesterol values were normalized to the internal standard and the mass of the sample.

### 4.10 High pressure freezing and electron microscopy

After sacrificing zebrafish, the spinal cord was dissected into three segments using the number of vertebrae for orientation. A 3mm long piece of the proximal and distal part of the spinal cord was high pressure frozen with the Leica HPM100 (Leica, Germany) using 20% polyvinylpyrrolidone (PVP, MW 10,000) (Sigma-Aldrich, Germany) in PBS as a cryoprotectant. The samples were then freeze-substituted using the Leica AFS2 (Leica, Germany) and embedded in Epon (Serva GmbH, Germany), as previously described (Weil et al., 2019).

Semithin (0.5 µm) or ultrathin (50 nm) sections of Epon embedded tissue were cut using the Leica UC7 ultramicrotome (Leica, Germany) and contrasted with UranylLess stain (Science Services GmbH, Germany). Electron micrographs were acquired using the electron microscope LEO EM912AB (Carl Zeiss Microscopy GmbH, Germany) equipped with a wide-angle dual speed 2k-charged-coupled device (CCD)-camera (TRS, Germany).

### 4.11 Micro-computed tomography (CT) scan

Zebrafish were sacrificed, briefly rinsed in water and then transferred to 35% and 70% ethanol for 1 hour each. Staining and fixation were carried out by placing fish for approximately 10 days at room temperature (RT) under slow rotation in a 4% PFA solution (Serva Electrophoresis, Germany) in PBS, pH 7.4 (Invitrogen, Germany), containing 0.7% phosphotungstic acid (PTA) solution (Merck (Sigma-Aldrich), Germany) diluted in 70% ethanol. Afterwards, fish were briefly rinsed in water and embedded in 1% agarose (Carl Roth, Germany) in 1.8 ml vials (Nunc CryoTube Vials, Merck (Sigma-Aldrich), Germany). For scanning, the in-vivo micro-CT system Quantum FX (Perkin Elmer, USA) was used with the following settings: tube voltage 90 kV, tube current 200 µA, field of view (FOV) 10×10 mm^2^, total acquisition time 3 min, resulting in a reconstructed pixel size of 20 µm and an image matrix of 512×512×512 voxel (Dullin et al., 2017). A custom made python script was used to stitch 2-3 consecutive scans together. This script, in addition to finding the perfect overlap also corrects for drift in between the scans. Data was analysed using NIH ImageJ/FIJI software (Schindelin et al., 2012).

### 4.12 Statistics

Statistical significance for all data sets presented in this study were calculated using Fisher’s permutation test (Crowley, 1992, Ernst, 2004, Fisher, 1970) except for mass spectrometry analysis of cholesterol which was calculated using Wilcoxon rank-sum (Mann and Whitney, 1947). For the permutation test, the original samples were bootstrapped 200,000 times. All p values were corrected with Benjamini-Hochbergs false discovery rate correction (Benjamini and Hochberg, 1995).

## 5 Results

### 5.1 Loss of Spastizin from the brain and spinal cord of *spastizin* mutants

In this study, we sought to uncover the role of SPASTIZIN in the pathogenesis of HSP, a debilitating disorder characterised by progressive lower limb spasticity and weakness. We previously established that “*souffle* ” is the zebrafish orthologue of the HSP-associated gene, *ZFYVE26* which encodes SPASTIZIN (Kanagaraj et al., 2014). Further analysis of the p96re allele of *souffle*/*suf* revealed that Spastizin lost only 282 of 2552 amino acids in the full-length protein and no significant differences in the locomotion of *suf* mutants were found when compared to the wild-types. We therefore generated a stronger mutant allele named “*zfyve26^ge1^*”. Molecular characterization of the mutated *zfyve26* gene revealed a four nucleotide deletion in the second exon, leading to a frameshift mutation that introduced a premature stop codon after amino acid 86 [Supplementary Figure 1 & Figure 1A]. Together, these results suggest that *zfyve26^ge1^* is a mutant allele with reduced Spastizin activity. To test this hypothesis, we conducted quantitative PCR experiments, which suggested elevated levels of mRNA in the brain and spinal cord of the mutants. For knockout zebrafish mutants, an increase in mRNA levels has been reported before (Rossi et al., 2015), indicating cellular mechanisms that compensate for the lack of protein in cells (El-Brolosy et al., 2019). Indeed, we found a significant decrease in the expression of Spastizin protein in the brain and spinal cord of the mutants, analysed using Western blot and immunohistochemistry, respectively. Whereas in Western blots a band of the expected size (>250 KDa) was visible in both wild-type and heterozygous mutant extracts, the bands intensity was significantly reduced for homozygous mutants [Figure 1B]. As a loading control, we used anti-*β*-Tubulin protein, for which a band of 55 KDa was observed in the brain tissue of each genotype [Supplementary Figure 3A]. To analyse the spatial expression of Spastizin, we examined sections of spinal cords with anti-Spastizin antibody. Remarkably, we observed high expression of Spastizin protein in the large axons of the M-cells of wild-type fish [Figure 1C & D, Supplementary Figure 3B]. Moreover, Spastizin expression was significantly reduced in the axons of the homozygous mutants. Quantitative analysis of the staining revealed a statistically significant reduction of Spastizin in homozygous mutants, further supporting the notion that Spastizin is not present in the neuronal tissue [Figure 1E]. Taken together, our results suggest that Spastizin protein is highly expressed in M-cells and this expression is lost in *zfyve26^ge1^* mutants.

**Figure 1:**
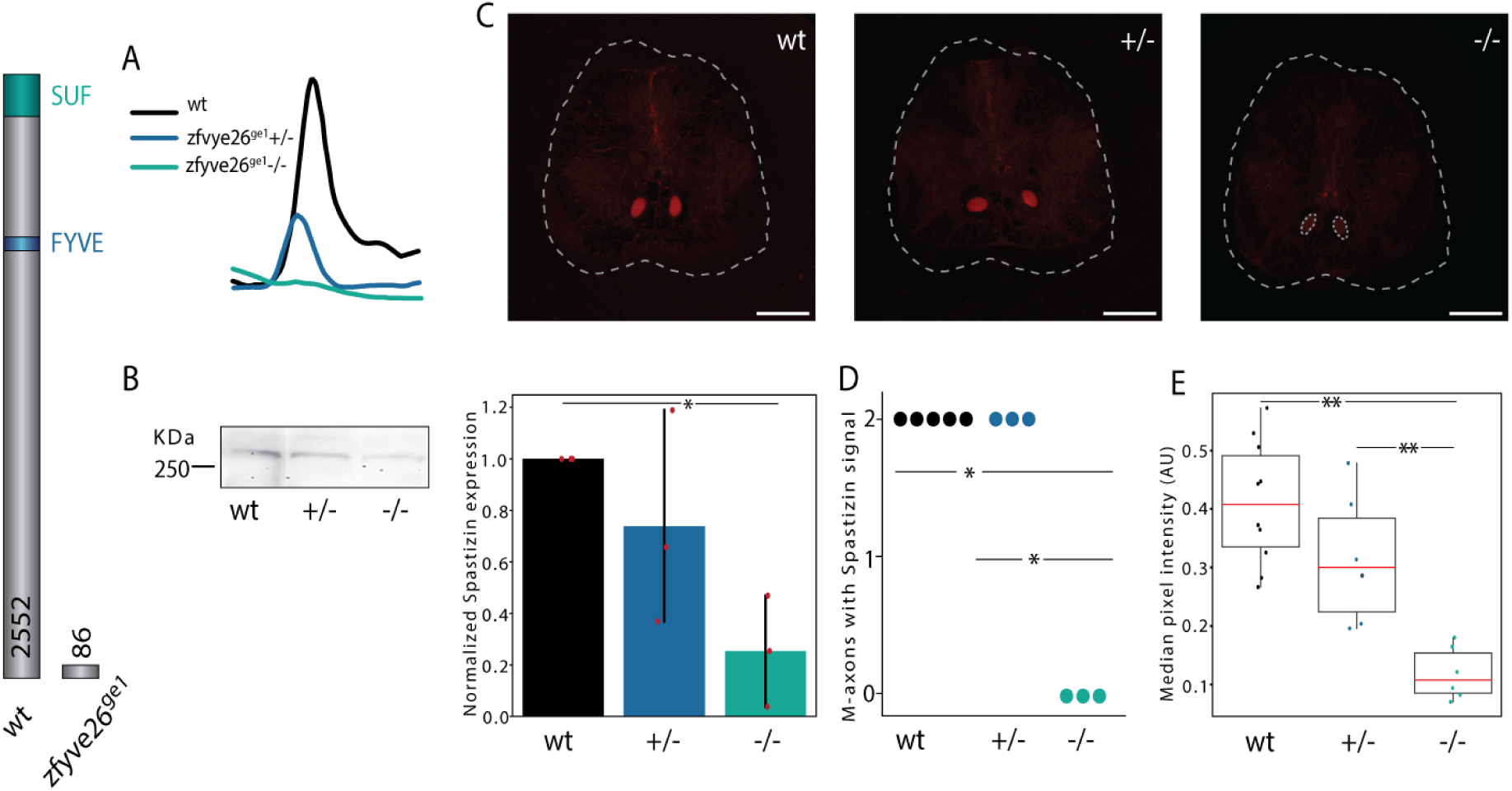
Effects of CRISPR-Cas9 mediated gene editing on Spastizin expression. A) A four basepair deletion in the second exon leads to the production of a truncated Spastizin protein of 86 amino acids. B) Immunoblotting of brain tissue, showing reduced levels of Spastizin protein in the homozygous mutants. Bar graph shows relative expression of Spastizin protein, normalized to anti-*β*-Tubulin, between the three genotypes. Data is depicted as mean and variance with red dots representing the individual data points. There is a significant decrease in the expression of Spastizin in the brain of mutant fish compared to the wild-type. (n, no. of experimental repeats = 3). C) Representative images of the immunohistochemical analysis of the sections of the spinal cord using anti-Spastizin primary antibody and donkey anti-rabbit Cyanine Dye 3 (Cy3) secondary antibody. The white dashed lines mark the section of the spinal cord. The red area illustrate the Mauthner axons with Spastizin expression. Although Spastizin was expressed in both the axons in wild-type and heterozygous mutant fish, it’s expression was significantly reduced in the axons in homozygous mutants, marked by the white-dashed circles (Scale bar: 100 µm). D) Quantification for the number of Mauthner axons having a signal for Spastizin in each genotype. The dots represent the number of animals examined from each genotype. E) Box plot of the median pixel intensity of the Spastizin signal in the two M-cell axons, normalised to the maximum intensity. The red line represent the median of all values, the box displays the upper and lower quartile, the whiskers denote 1.5 times the interquartile range and the dots show individual data points. There is a significant reduction of the signal intensity for Spastizin in the M-cell axons of homozygous mutants compared to both heterozygous and wild-type fish. (N, no of fish, wt = 5, +/- = 3, -/- = 3). Statistical significance was tested with independent t-test for Spastizin expression and by Fisher’s permutation test for number of M-axons with Spastizin signal and its intensity in the M-axons. *p *<* 0.05, **p *<* 0.01.

### 5.2 Locomotion defects in *spastizin* mutants

To address whether Spastizin is required for locomotion in zebrafish, we first tested the ability of the animals to swim against a water stream that induced their natural rheotaxic behaviour (Arnold, 1974). In *spastizin* mutants, we observed a significantly reduced ability to remain in the centre of the stream [Figure 2A]. Using 2-dimensional (2-D) heat maps of the probability density for every possible position along the setup, we found that while wild-type zebrafish preferred to remain in the centre of the current, heterozygous and homozygous mutants avoided it [Figure 2B]. Apparently, *spastizin* mutants have less endurance than wild-types in swimming constantly against the water stream. Zebrafish create translational thrust through undulatory sinus-like waves of their body. This motion is composed of a standing and a travelling wave (Gray, 1933, Lauder and Tytell, 2005, Muller et al., 2000, Müller and Van Leeuwen, 2006, Smits, 2019, Wardle et al., 1995). Wild-type zebrafish create both waves during movement [Figure 2F, full flexibility wave], but our data reveals that this travelling wave is dampened at the caudal end of *spastizin* mutants [Figure 2F, stiff wave]. While a complete loss of the abdominal travelling wave would lead to a severe reduction of the thrust speed, it only slightly decreased in the mutants [Figure 2C]. Possibly, the mutant fish compensate for their caudal spasticity, potentially leading to a reduction in propulsion efficiency that prevents them from reaching the same rheotactic endurance as wild-type fish. Indeed, our analysis revealed that the mutants bend their body more strongly to achieve the same acceleration as the wild-type fish [Figure 2D], suggesting that the increased pectoral standing wave might be compensating for the loss of the travelling wave. To substantiate this interpretation, we developed a computational model of the propulsion with either a reduced travelling wave (stiffness), or with the reduced travelling wave being supplemented by an increased standing wave (pectoral fin). This confirmed that a standing wave of the same angle but with a higher amplitude can compensate for the loss of the travelling wave in the mutants [Figure 2F, compensatory wave]. For this, the peak of the standing wave would have to travel more caudally [Figure 2F]. Indeed the data of heterozygous and homozygous mutants showed that the peak of the pectoral bending shifts more caudally as compared to the wild-types [Figure 2E]. These results suggest that we established the first genetic HSP model in adult zebrafish with locomotion defects.

**Figure 2:**
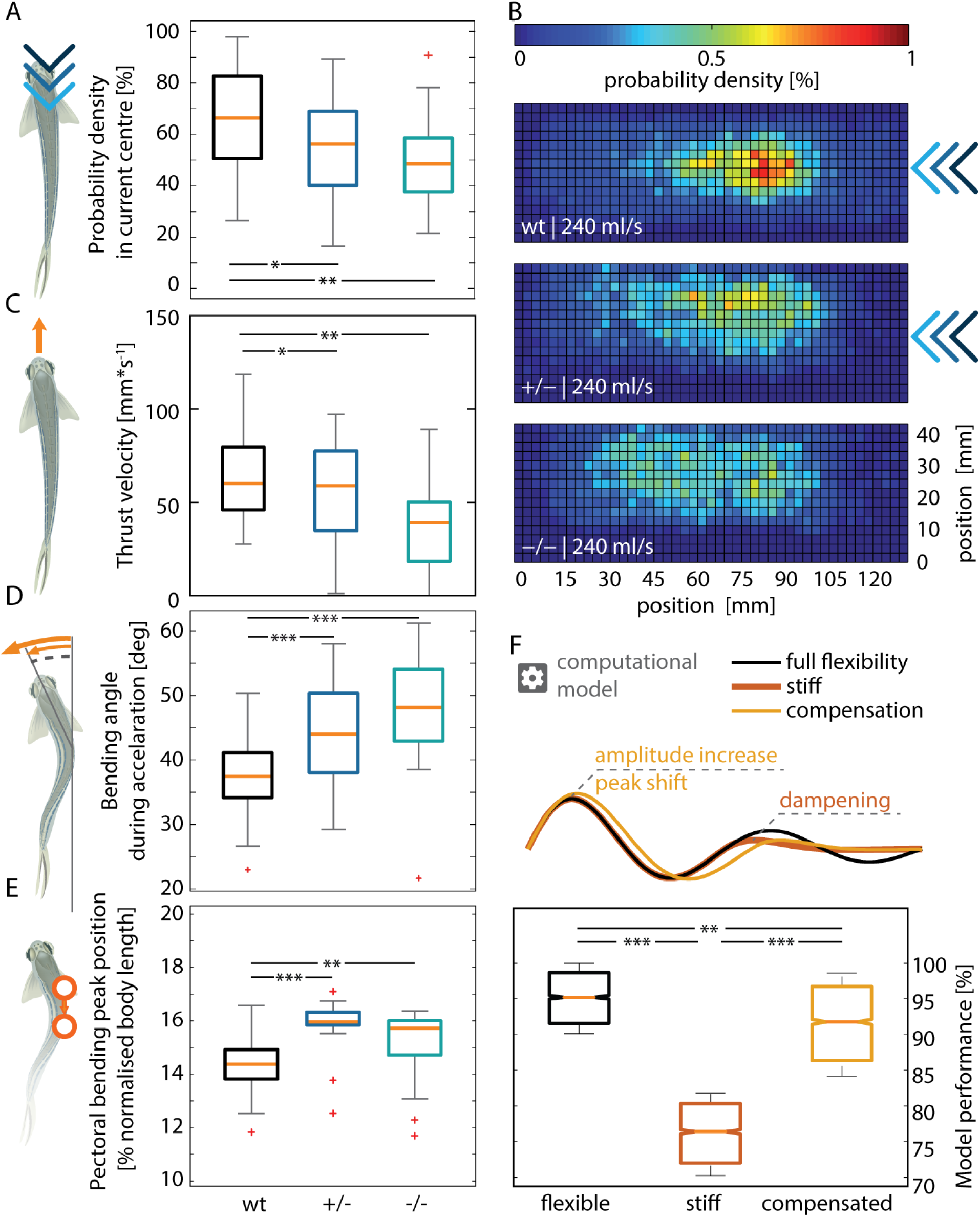
Locomotion defects observed in *spastizin* mutants during unmotivated and counter current trials. Data is represented as box plots for assessing various aspects of locomotive ability of wild-type and *spastizin* mutants. The orange line shows the median of all individuals, the box displays the upper and lower quartile, the whiskers denote 1.5 times the interquartile range, the plus-sign marks outlier. A) The homozygous mutants spend less time in the centre of the water current as compared to heterozygous and wild-type fish. B) Each histogram is normalized to 100% and the color bar goes from 0 to 1%. The heat map shows all possible locations of the fish in the setup. Blue colour indicates low probability and red colour illustrate high probability of presence. Whereas the *spastizin* mutants are more dispersed within the setup, the wild-type fish stay more at the centre, where the current is most constant. C) Decrease in the median thrust velocity of homozygous *spastizin* mutants as compared to heterozygous and wild-type fish during unmotivated trials. D) Increase in the bending angle for both heterozygous and homozygous mutants compared to wild-types. E) Increase in pectoral bending peak position (normalised to body length) of heterozygous and homozygous mutants compared to wild-types. F) Computational model depicting a shift in the peak position of pectoral standing wave which compensates for the dampened travelling wave. The box plot shows the model performance with fully flexible and compensated at the same level and high as compared to the stiff. (N, no. of fish, wt = 53, +/- = 23, -/- = 21). Statistical significance was tested with Fisher’s permutation test. *p *<* 0.05, **p *<* 0.01, ***p *<* 0.001.

### 5.3 Functioning of M-cells affected in *spastizin* mutants

M-cells drive the C-start escape response (Eaton et al., 1977, Faber and Korn, 1978). Because these cells are giant neurons, their field potential can be measured in freely swimming fish using the electrodes attached to the fish tank (Issa et al., 2011). We here exploit this to investigate if the lack of Spastizin from the M-cells resulted in altered escape behaviour and/or could be linked to altered M-cell activity in evoked escape responses. Escape behaviour in zebrafish was elicited by administering an air puff into the tank using a micro spritzer pump.

Contrary to wild type fish, the mutants could not form a C-bend in response to the stimulus, as can be seen by the turtuosity of their silhouette ( *<u>^HeadT^ ^ailDistance−bodylength^</u>* , Figure 3A). Since the C-bend is crucial for the rapid acceleration during the escape response, mutants exhibit a discernibly slower movement compared to the wildtype (Figure 3B). In addition, we find that while both the small and large spike frequency was lower in spastizin mutants (Figure 3C, D, & Supplementary Figure 4), only large spikes occurred coupled to the stimulus (time 0). These observations document that Spastizin has an influence on the activity of the M-cells, with its significant reduction contributing to a distinctly slower escape response.

**Figure 3:**
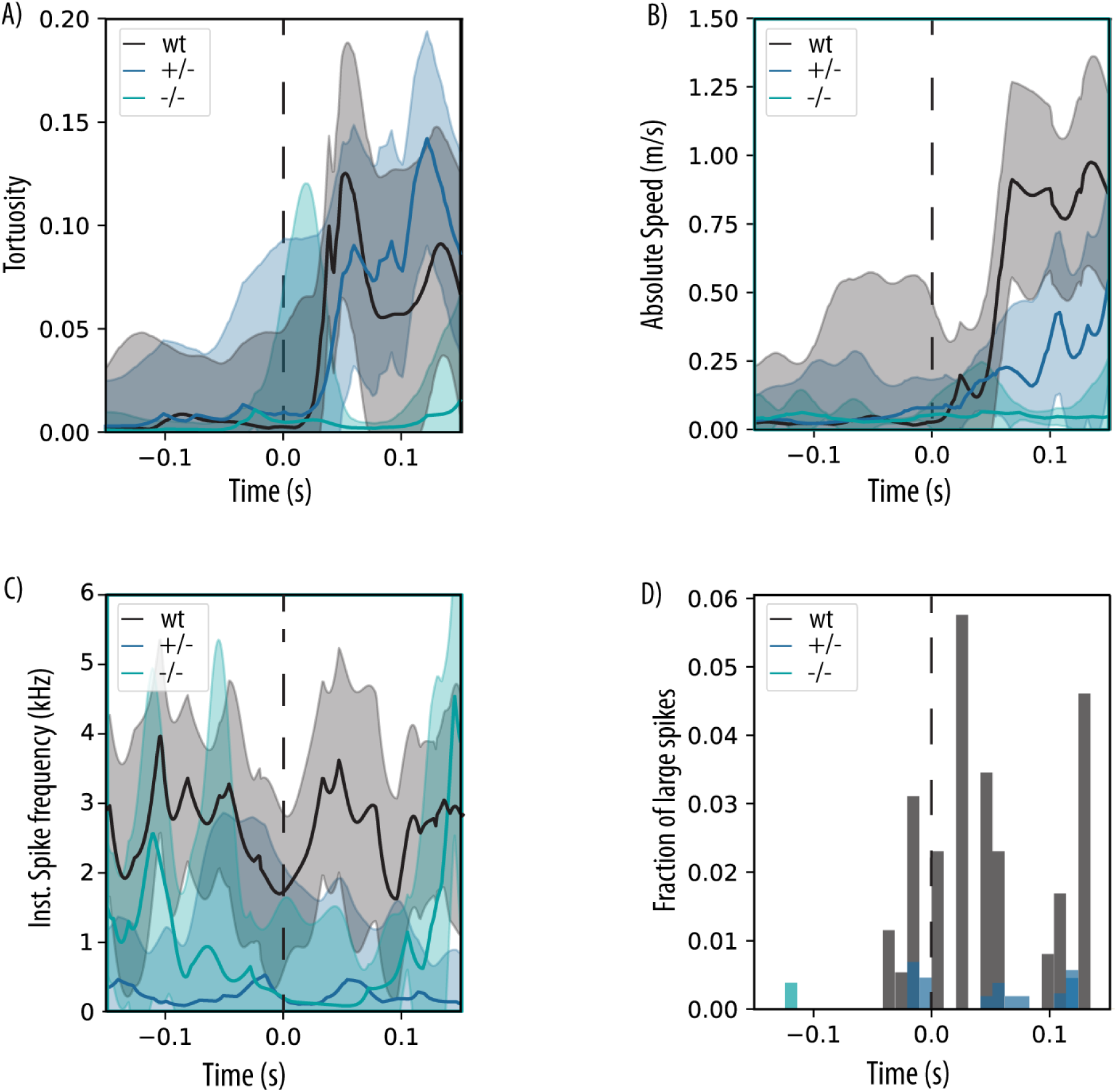
Lack of Spastizin affects the function of M-cells. A) shows the tortuosity of the zebrafish over time as median with the surrounding 95% confidence interval. Tortuosity is defined as the difference of the body length and the distance between the head and tail of the animal, normalised to the body length. A value of 0 signifies the animal is entirely straight and 1 tortuosity signifies that head and tail touch. B) presents the absolute velocity during the same time window. C) shows the instantaneous spike frequency over time, as in A. The dashed vertical line marks the time of the stimulus presentation. D) shows the mean histogram of the occurrence of large spikes during the same time window. All the three parameters are low for homozygous mutants compared to heterozygous and wild-types. (N, no. of fish, wt = 9, +/- = 18, -/- = 5).

### 5.4 Degeneration of lower motor neuron axons and their myelin sheaths in the *spastizin* mutants

Axon degeneration of the corticospinal motor neurons in a length dependent manner is the prime cause for spasticity and locomotion defects in HSP patients (Blackstone, 2012, 2018, Finsterer et al., 2012, Klebe et al., 2015, Tesson et al., 2015). Interestingly, we also found a significant decrease in the number and diameter of axons in the spinal cord of the *spastizin* mutants [Figure 4A, B, & C], confirming axonopathy in the zebrafish model of SPG15. We also found a significant decrease in the abundance of cholesterol in the brain of *spastizin* mutants [Figure 4D], and we thus investigated the cellular structure of the neurons in the spinal cord. Compared to the intact myelin observed in wild-types [Figure 4E, F, G, & H], severe splitting and vesiculation, two hallmarks of demyelination, were present in the large-calibre axons of mutant spinal cord motor neurons [Figure 4I, J, & K]. Similar features of myelin sheath degeneration have been observed in rat models of demyelinating diseases such as neuromyelitis optica (NMO) and experimental autoimmune encephalomyelitis (EAE) (Weil et al., 2016). In addition to demyelination, we found axonal swelling in the *spastizin* mutants [Figure 4L], a further distinctive feature of neuropathology. Our results thus indicate that Spastizin is essential in the maintenance of the myelin sheath and axonal integrity. Together, these results suggest that the locomotion defects are caused by demyelination and degeneration of the lower motor neurons in the *spastizin* mutants.

**Figure 4:**
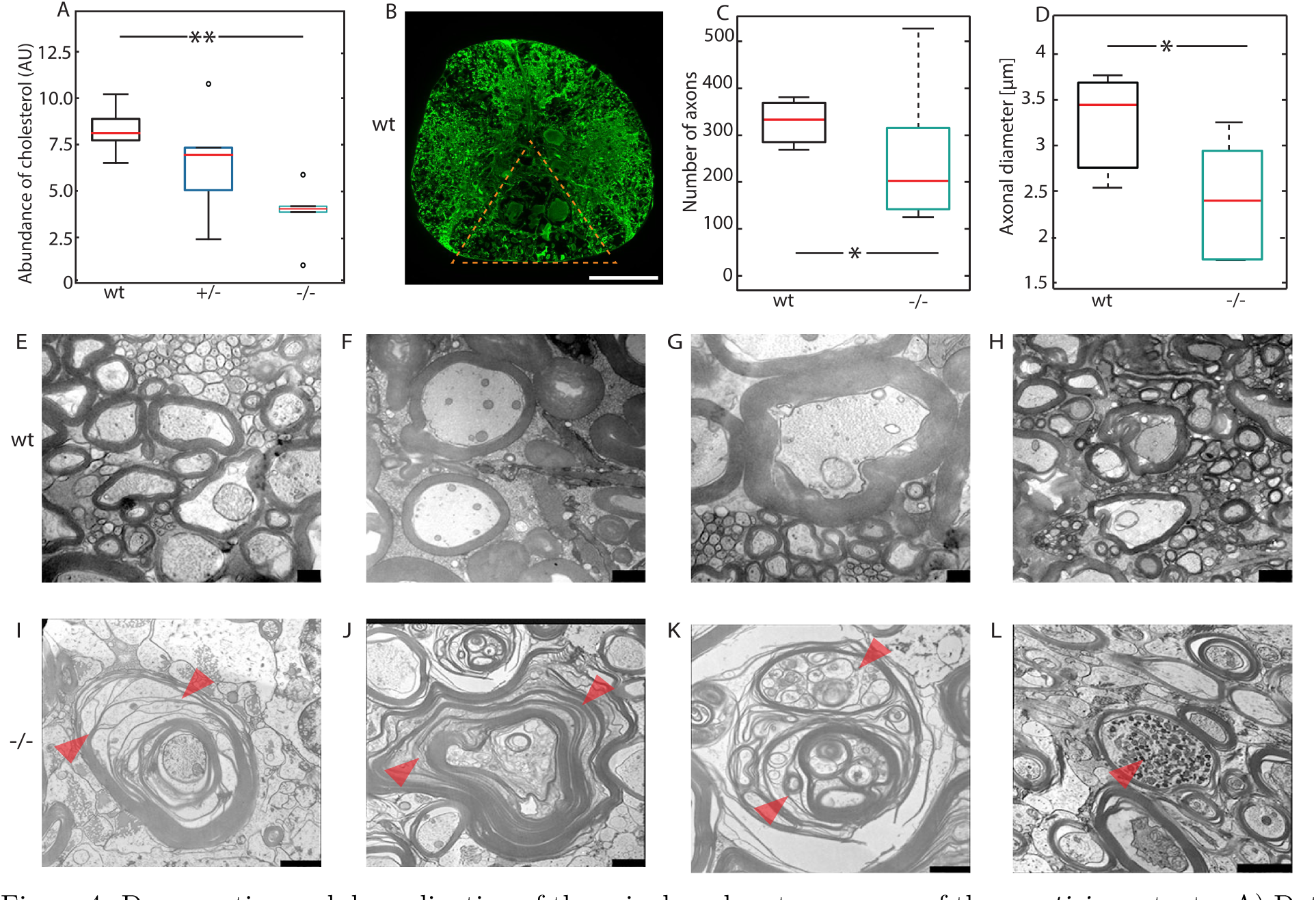
Degeneration and demyelination of the spinal cord motor neurons of the *spastizin* mutants. A) Data is represented as box plots for the abundance of cholesterol in the brain of wild-type and *spastizin* mutant fish, divided by the mass of the sample and normalized to the internal standard. The red line depicts the median, the box displays the upper and lower quartile and the whiskers denote 1.5 times the inter quartile distance. Black circle marks outlier. There is a significant reduction in the abundance of cholesterol in the brain of homozygous mutants compared to the wild-types (N, no. of fish, wt = 5, +/- = 5, -/- = 5). B) Representative image of immunohistochemical analysis of spinal cord cross sections stained with a neuronal marker anti-*β*-Tubulin. The triangle marks the area in which the neurons were quantified (Scale bar: 100 µm). C), & D) Data is represented as box plots for number and diameter of axons in the evenly distributed sections of the spinal cord for wild-type and mutant zebrafish. The red line depicts the median, the box displays the upper and lower quartile and the whiskers denote 1.5 times the inter quartile distance. There is a significant decrease in the number and diameter of axons in the *spastizin* mutants compared to the wild-types (N, no. of fish, wt = 3, -/- = 6). E)- L) Representative images of the cross section of spinal cord showing large caliber axons surrounded with myelin sheath. E), F), G) & H) Representative images of intact myelin sheath in the wild-type fish. I) & G) Arrow heads mark fragmentation or splitting of myelin sheath in the homozygous mutants. K) Severe vesiculation profile of myelin sheath marked by arrow heads, in homozygous mutants. L) Arrow head indicates swelling in large caliber axons of the spinal cord of homozygous *spastizin* mutants. (Scale bars: 250 nm in E) & G), 1 µm in F), H), I), & K) and 2 µm in J) & L)). Statistical significance was tested with Wilcoxon rank-sum test for cholesterol abundance and with Fisher’s permutation test for the number and diameter of axons *p *<* 0.05 and **p *<* 0.01.

### 5.5 No skeletal deformities in *spastizin* mutants

To further validate that the locomotion defects in *spastizin* mutants are caused by neuronal defects rather than skeletal abnormalities, we utilised micro-computed tomography (CT) scanning to analyse the vertebral column of wild-type and *spastizin* mutants. Our analysis revealed no distinguishable differences in the skeletal structure between wild-type and heterozygous or homozygous mutants [Supplementary Figure 5]. In addition, we examined the muscle area in the caudal region of the fish. It was unaltered in the mutant fish [Supplementary Figure 6], indicating absence of muscular atrophy. Hence, instead of reporting muscle defects, the observed behavioral phenotypes seem to be caused by neuronal defects.

## 6 Discussion

Hereditary spastic paraplegia (HSP) is a complex disorder associated with more than 88 genes and several cellular pathways (Blackstone, 2012, 2018, D’Amore et al., 2018, Elsayed et al., 2021, Fink, 2014, Galatolo et al., 2018, Giudice et al., 2014, Hensiek et al., 2015, Parodi et al., 2018). The respective pathophysiological mechanisms, however, are little understood, hampering therapeutic treatments of HSP (Meyyazhagan and Orlacchio, 2022, Shribman et al., 2019). Using zebrafish as a model system, we found that Spastizin protein is particularly abundant in the large-caliber axons of the Mauthner-cells, and that its reduction causes degeneration of the myelin sheath and locomotion defects, recapitulating key disease features of HSP. Studies in zebrafish oocytes, mouse models, and patient-derived cells have suggested that Spastizin plays a crucial role in various cellular processes, such as membrane trafficking (Kanagaraj et al., 2014, Khundadze et al., 2013), autophagosome maturation (Vantaggiato et al., 2013, 2019), lysosomal regeneration (Chang et al., 2014), cytokinesis (Sagona et al., 2010), which all seem particularly important for the survival and maintenance of large neurons (Blackstone et al., 2011, De Vos and Hafezparast, 2017, Stavoe and Holzbaur, 2019). Through which of these mechanisms, Spastizin causes demyelination and cell degeneration can now be studied using Mauthner cells, whose behavioural relevance and physiology have been investigated extensively (Korn and Faber, 2005).

The observed dysfunction in Mauthner cells parallels the loss of limb movement control in human patients (Blackstone, 2012, 2018, Fink, 2006, 2014, Finsterer et al., 2012, Harding, 1993, 1984, Klebe et al., 2015, Tesson et al., 2015). Frogs, uniquely possessing both Mauthner cells and legs, provide a critical model for our understanding: Post-metamorphosis, these cells in frogs are known to control limb movements (Davis Jr and Farel, 1990, Will, 1986), thereby bolstering the argument that Mauthner cells offer a viable model for studying lower limb control issues characteristic of HSP. The study also noted an impact on other large motor neurons, contributing to reduced mobility during normal cruising behaviour. This reduction, however, is often masked by compensatory movements in the fish, potentially explaining the limited research on adult fish in the context of motor neuron degeneration. Despite such compensation mechanisms, a notable decrease in thrust velocity was observed in the mutants. To ascertain that this decrease stems from motor neuron dysfunction rather than skeletal issues, CT scans were conducted. We found no significant differences in the vertebral column or muscle mass between wild-type and *spastizin* mutants, confirming that the locomotion defects arise from motor neuron degeneration rather than skeletal deformities.

Recent studies implicate altered lipid metabolism, particularly cholesterol, in HSP pathophysiology (Dai et al., 2021, Darios et al., 2020, Mou et al., 2020, Rickman et al., 2020, Vance, 2012). Cholesterol, vital for membrane and myelin formation (Krause and Regen, 2014, Saher et al., 2005, 2011, Zhang and Liu, 2015), is affected in *spastizin*mutants, with reduced brain cholesterol levels hinting at disrupted regulatory mechanisms. This aligns with Spastizin’s localisation at the ER, potentially influencing cholesterol metabolism (Hanein et al., 2008, Murmu et al., 2011). Furthermore, our findings of demyelination and neuropathology in *spastizin* mutants spinal cords suggest a novel role for Spastizin in myelin integrity, potentially linked to cholesterol abundance. The interplay of Spastizin with calcium homeostasis, akin to its interaction partner Spatacsin’s effect on calcium and cholesterol levels (Boutry et al., 2019), warrants further investigation to elucidate its role in HSP pathogenesis.

## 7 Conclusions

Overall, our study provides further insights into the role of Spastizin in the pathogenesis of SPG15. Our study suggests that depreciation of Spastizin from the large caliber axons disturbs the metabolism of an essential component of myelin which affects its integrity, causing large caliber axons to degenerate. This leads to weakness, spasticity, and other locomotion problems, particularly defects in the M-cells. As electron microscopy analysis, biochemistry and measuring the activity of M-cells in a fully functional organism is not possible in other vertebrates, our study opens novel avenues to address the cellular and biochemical processes affected in HSP and establishes a novel tool to develop therapeutic strategies for the benefit of HSP patients.

## 9 Ethics Statement

The zebrafish were maintained according to the guidelines provided by the Westerfeld zebrafish book (Westerfield, 2000) and EuFishBioMed/Felasa (Aleström et al., 2020), in compliance with the regulations of the Georg-August University, Goettingen, and Bielefeld University (540.4/16 25/Engelmann), Bielefeld, Germany. The zebrafish experiments were approved by the Lower Saxony State Office for Consumer Protection and Food Safety (AZ14/1681/Dosch) and were performed according to EU directive 2010/63/EU.

## 10 Consent to participate

Not applicable

## 11 Consent for publication

Not applicable

## 12 Data Availability Statement

The data generated for this study is included in the article and supplementary material and will be available on reasonable request by the corresponding author.

## 15 Authors Contribution

BG, VG, and RD designed the study. VG, SA, and LH performed experiments and data analysis. VG, PS, TI performed data analysis for cholesterol abundance. GK generated the mutant line. WM & TR generated EM data. CD generated CT scan data. VG and BG wrote the first draft of the manuscript with editing and inputs from all authors.

## 16 Acknowledgements

We would like to thank Nicola Schwedhelm-Domeyer, Stephanie Pauls, Silvia Gubert, and Tatjana Openkowski for technical assistance and Gudrun Kracht, Axel Zigan, and Helene Schellenberg for animal care.

## 17 Supplementary material

**Figure 1:**
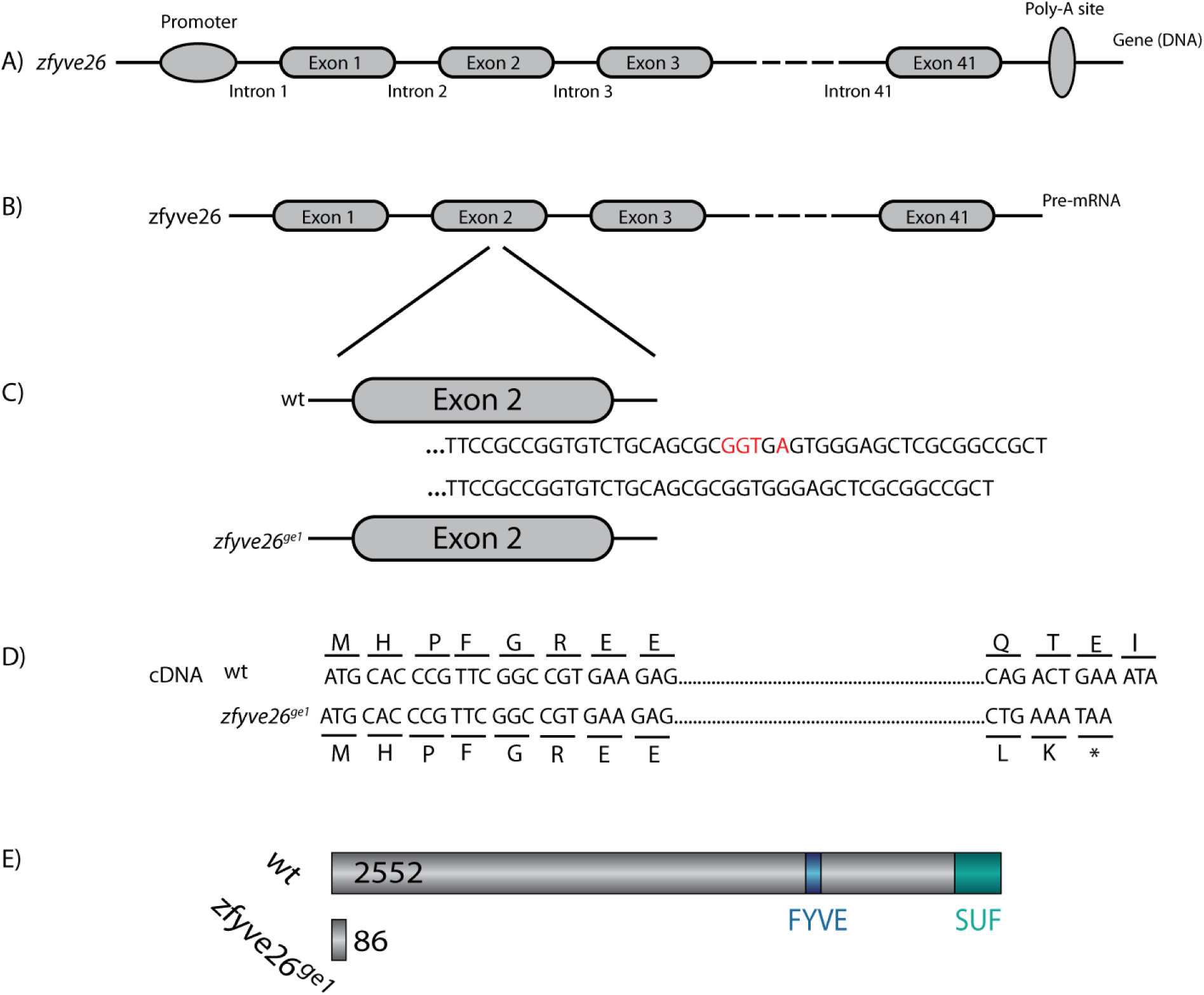
Descriptive representation for the generation of *zfyve26^ge1^* allele. A) & B) DNA and pre-mRNA structure of *zfyve26* gene. C) & D) Deletion of four bases highlighted in red color from exon 2 leads to a frameshift mutation that eventually introduces a premature stop codon after amino acid 86. E) Representative picture highlights the structural differences between wild-type and the mutant protein. While wild-type protein is full in length with all essential domains, the mutant protein is only 86 amino acid long.

**Figure 2:**
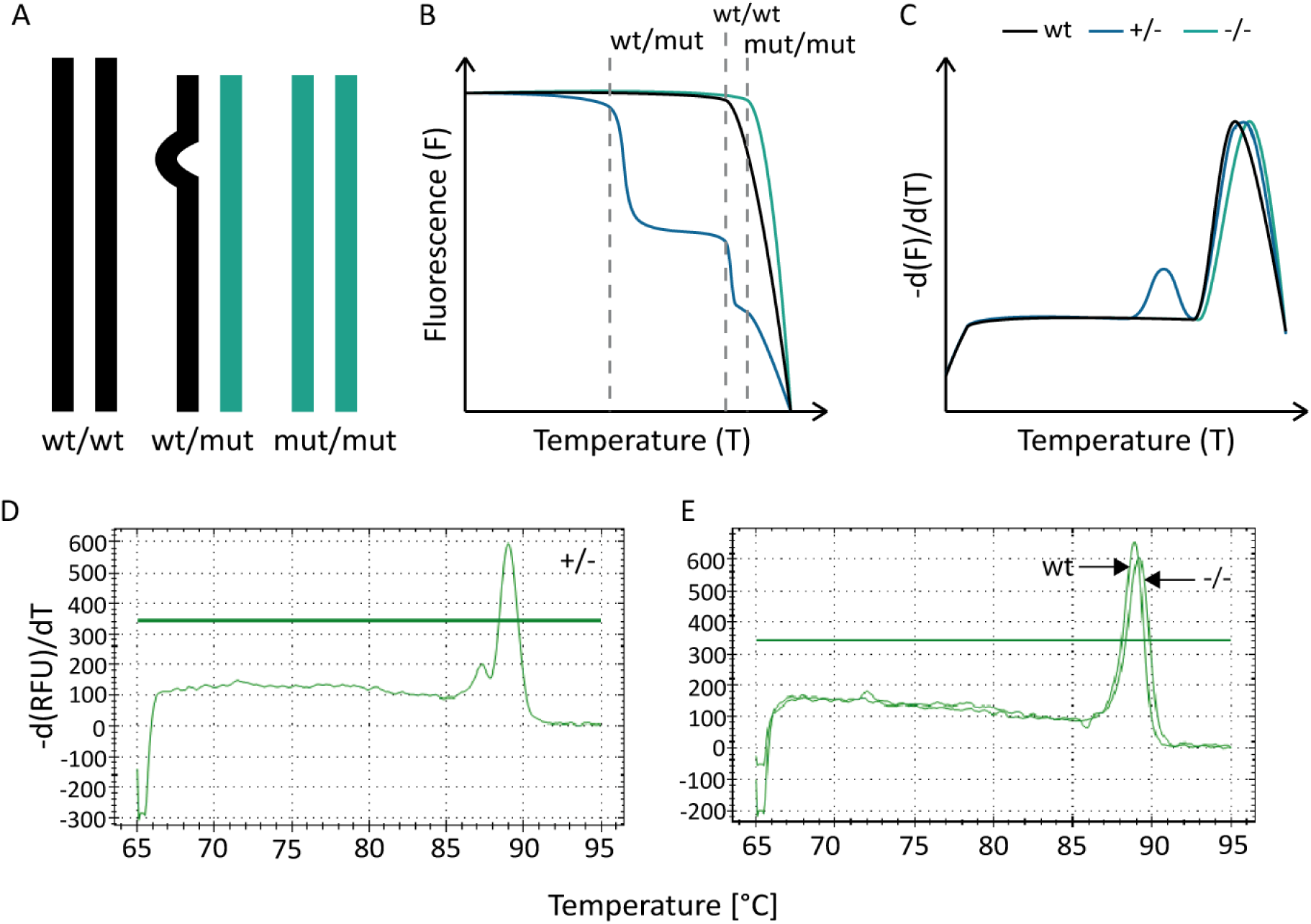
Melting curve analysis. A) Schematic of the three rehybridization products wt/wt, wt/mut, and mut/mut. B) Schematic of the expected melting curve for wild-type, heterozygous and homozygous mutant fish. Gray dashed bars indicate the relative melting points of each rehybridization product. As wt/mut is the most unstable product, it will dissociate first, resulting in a decrease in fluorescence intensity in heterozygous mutants. On the other hand, the melting temperature for mut/mut is slightly higher as compared to wt/wt. C) First derivate of B), in which peaks are better visualized. D) & E) Exemplary pictures of melt curve peaks for all the three genotypes. The heterozygous mutants are recognized by their shape which is distinguishable compared to homozygous mutants and wild-types. While the homozygous mutants are characterized by the temperature shift. The distinct peaks for homozygous mutants and wild-types are marked by arrows.

**Figure 3:**
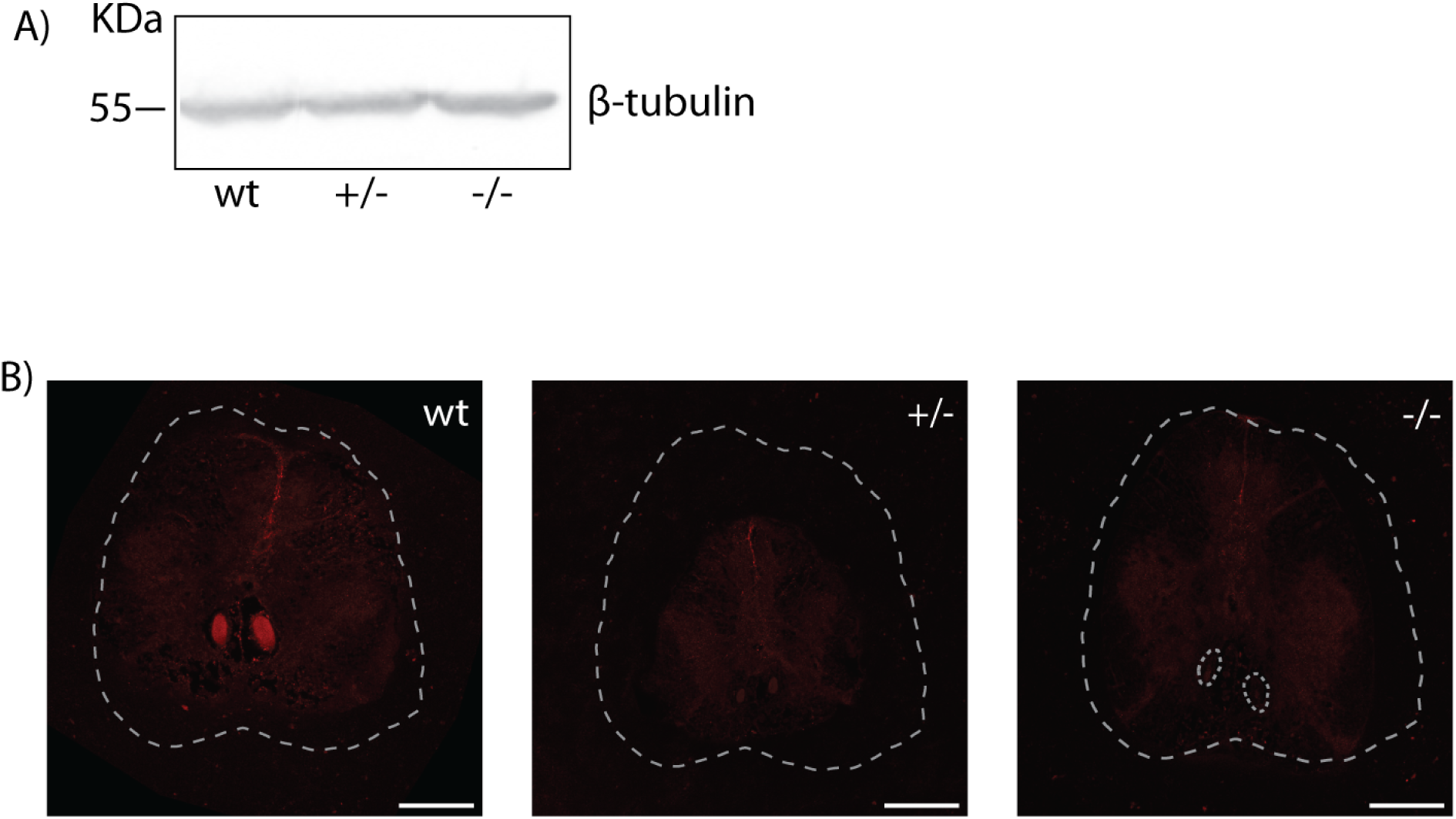
A) Immunoblot showing expression of *β*-Tubulin in the brain of wild-type, heterozygous, and homozygous *spastizin* mutants. B) Representative images of the immunohistochemical analysis of the sections of spinal cord using anti-Spastizin primary antibody and donkey anti-rabbit AF 488 secondary antibody. The white dashed lines mark the section of the spinal cord. The red area illustrate the Mauthner axons with Spastizin expression. Although Spastizin was expressed in both the axons in wild-type and heterozygous mutant fish, it was significantly reduced in the axons in homozygous mutants, marked by the white-dashed circles. (Scale bar: 100 µm).

**Figure 4:**
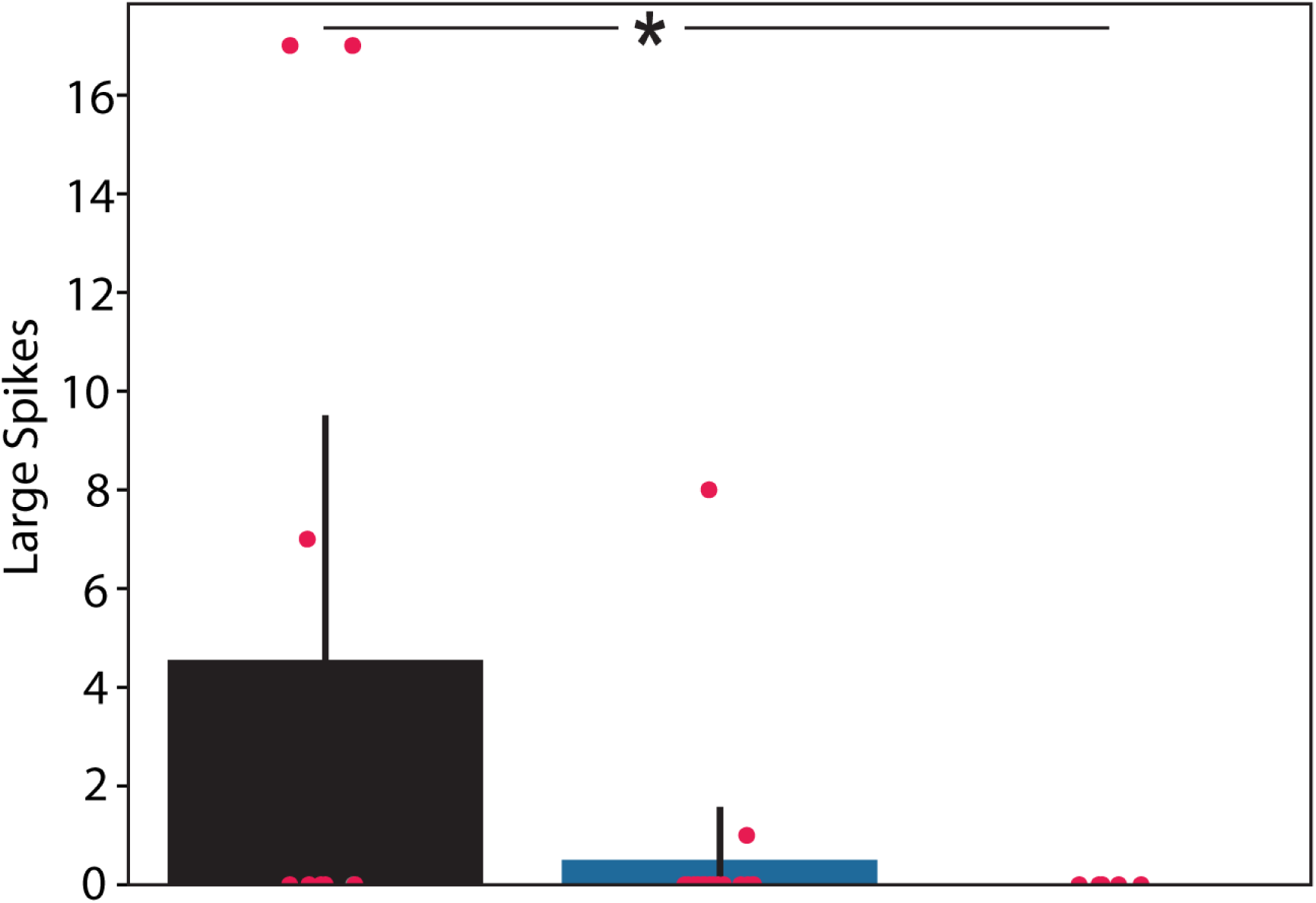
Effect of the reduction of Spastizin on the giant spikes. Bar graph shows the number of large spikes observed in the first 150ms post stimulus for the three genotypes. There is a significant reduction in the number of large spikes in homozygous *spastizin* mutants. Data is represented as mean *±* 95% confidence interval with red dots representing the individual data points. (N, no of fish, wt = 9, +/- = 18 , -/- = 5). Statistical significance was tested with Fisher’s permutation test. *p *<* 0.05.

**Figure 5:**
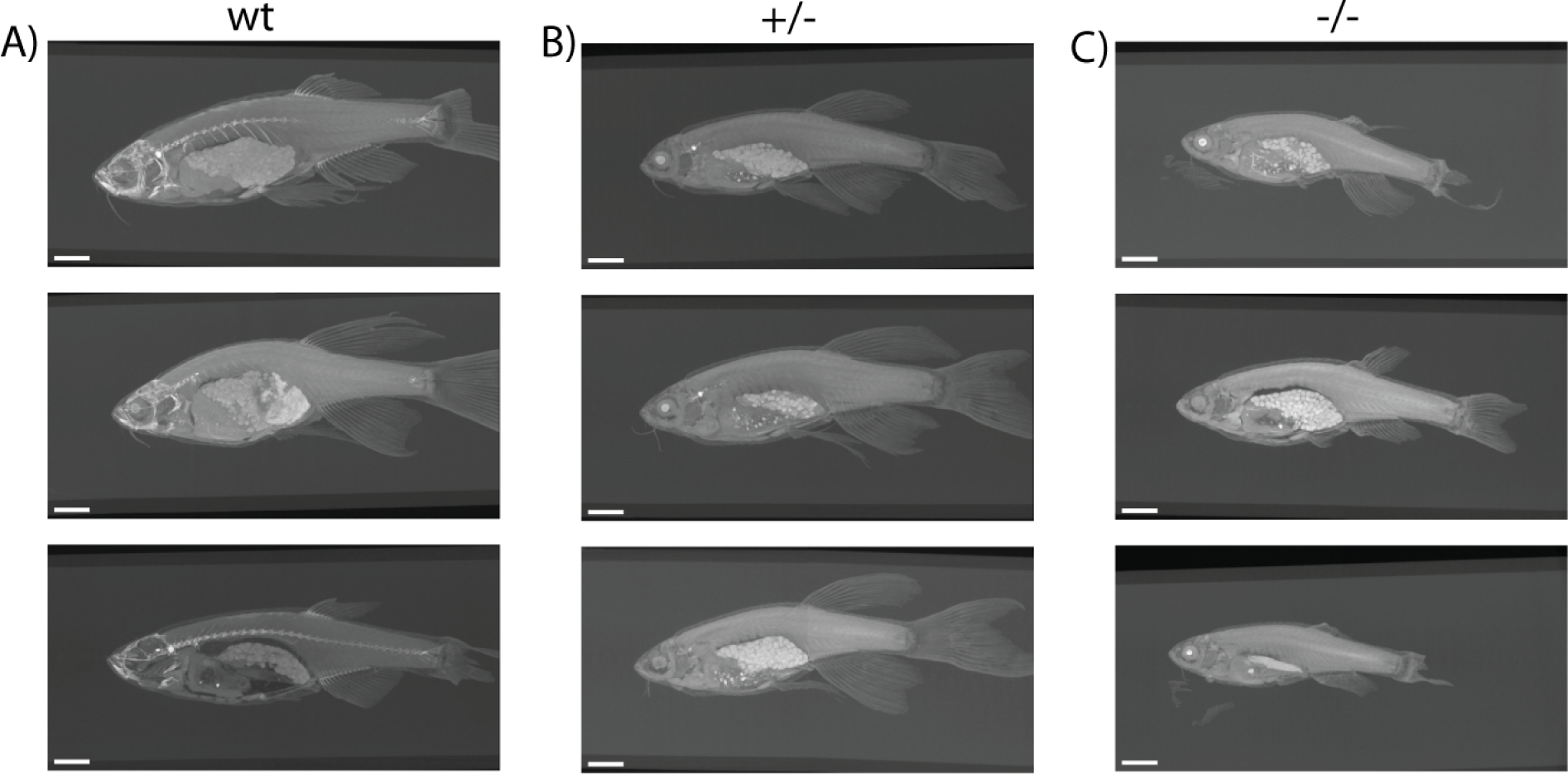
No skeletal changes in *spastizin* mutants. A), B), & C) Computed tomography images of the vertebral column of wild-type, heterozygous, and homozygous *spastizin* mutants. There is no change in the skeletal structure among the three genotypes. (Scale bar: 2 mm). (N, no. of fish analysed, wt = 3, +/- = 3, -/- = 3).

**Figure 6:**
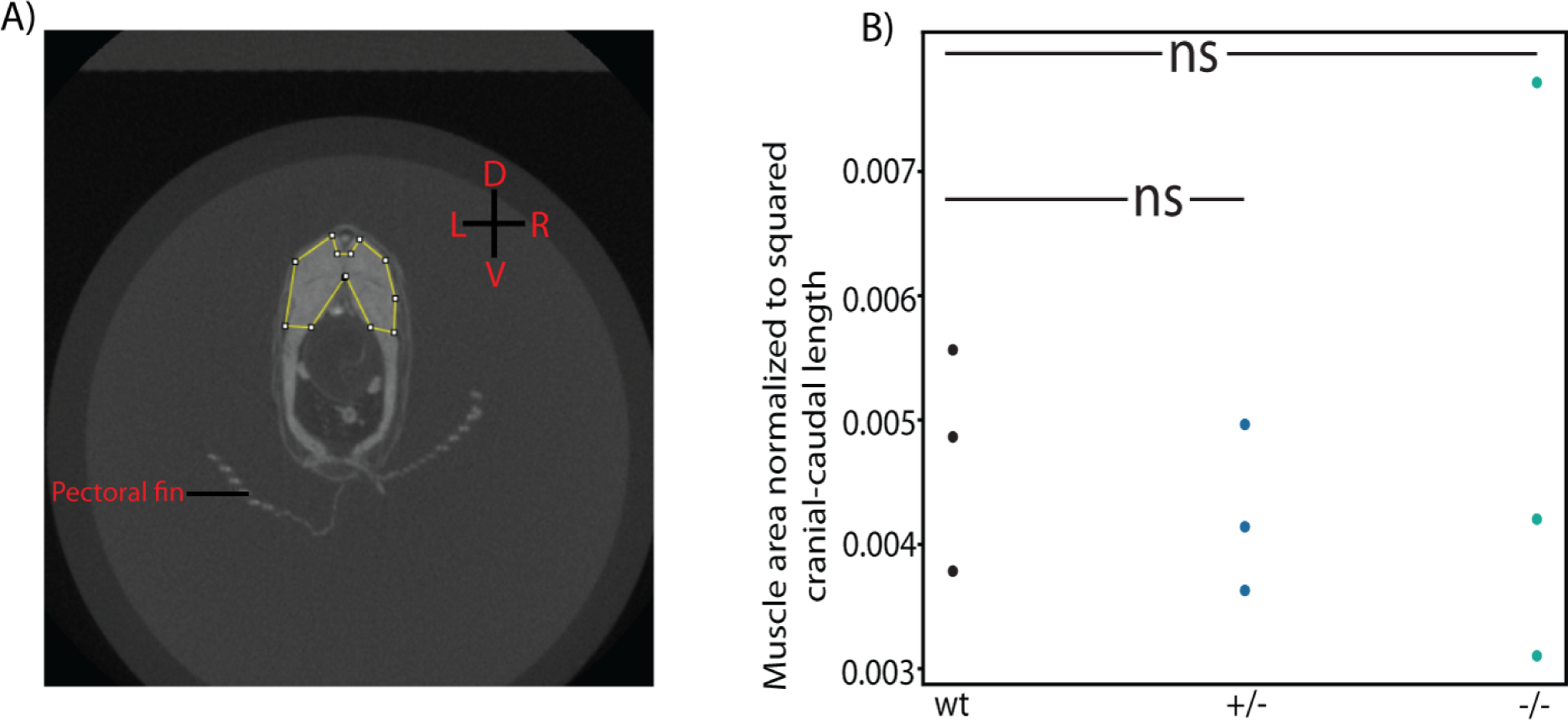
No muscle atrophy in *spastizin* mutants. A) Exemplary illustration of the fish body in transverse plane with the caudal muscles marked with the yellow segmented line. (D: Dorsal axis, V: Ventral axis, L: Left axis, R: Right axis). B) Scatter plot of caudal muscle area normalised to squared length of the fish body. There is no change in the muscle area in heterozygous and homozygous mutants compared to the wild-types. (N, no. of fish analysed, wt = 3, +/- = 3, -/- = 3) Statistical significance was tested using Fisher’s permutation test, ns is non-significant.

## 6 Abbreviations

HSP: Hereditary spastic paraplegia
SPG15: Spastic paraplegia 15
M-cell: Mauthner cell
ER: Endoplasmic reticulum
T7E1: T7 Endonuclease 1
FPS: Frames per second
LACE: Limbless animal traCkEr
GC-MS: Gas chromatography-mass spectrometry
CT: Computed tomography
2-D: 2-Dimensional
NMO: Neuromyelitis optica
EAE: Experimental autoimmune encephalomyelitis

